# In vitro biosynthesis of poly-β-1,4-glucan derivatives using a pro-miscuous glycosyltransferase

**DOI:** 10.1101/2020.02.14.949545

**Authors:** Gregory S. Bulmer, Ashley P. Mattey, Fabio Parmeggiani, Ryan Williams, Helene Ledru, Andrea Marchesi, Lisa S. Seibt, Peter Both, Kun Huang, M. Carmen Galan, Sabine L. Flitsch, Anthony P. Green, Jolanda M. van Munster

## Abstract

The β-1,4-glucose linkage of cellulose is the most abundant polymeric linkage on earth and as such is of considerable interest in biology and biotechnology. It remains challenging to synthesize this linkage in vitro due to a lack of suitable biocatalysts; the natural cellulose biosynthetic machinery is a membrane-associated complex with processive activity that cannot be easily manipulated to synthesize tailor-made oligosaccharides and their derivatives. Here we identify a promiscuous activity of a soluble recombinant biocatalyst, *Neisseria meningitidis* glycosyltransferase LgtB, suitable for the polymerization of glucose from UDP-glucose via the generation of β-1,4-glycosidic linkages. We employed LgtB to synthesize natural and derivatized cello-oligosaccharides and we demonstrate how LgtB can be incorporated in biocatalytic cascades and chemo-enzymatic strategies to synthesize cello-oligosaccharides with tailored functionalities. We also show how the resulting glycan structures can be applied as chemical probes to report on activity and selectivity of plant cell wall degrading enzymes, including lytic polysaccharide monooxygenases. We anticipate that this biocatalytic approach to derivatized cello-oligosaccharides via glucose polymerization will open up new applications in biology and nanobiotechnology.

## Introduction

Cellulose, a linear polysaccharide consisting of β-1,4-linked glucose, is of critical importance in biotechnology, nutrition and microbial pathogenicity. As major structural component in plant cell walls, it is the predominant constituent of lignocellulosic waste, a highly abundant (2 × 10^11^ tons per annum), energy-rich material that is exploited as a renewable resource in the bio-economy^1^. Cellulose conversion to sugar monomers enables biotechnology for sustainable production of fuels and chemicals^2^. Hydrolytic enzymes are deployed to this aim in commercial-scale biorefineries producing lignocellulosic ethanol and identification of effective enzymes attracts much interest^3–6^. Lignocellulose-derived cellulose and its oligosaccharides also function as dietary fiber, whereby their fermentation by intestinal microbiota produces short-chain fatty acids that promote gut health^7–9^. Cellulose is a structural component of oomycete cell walls^10^ and a potential target for their biocontrol^11^; it is critical for pathogenicity of the causal agent of potato blight, a devastating crop disease costing $5 billion per annum^12^. In bacterial biofilms associated with hard to eradicate, debilitating infections, cellulose functions as an essential matrix component, which is exploited as a biomarker for their detection^13–19^. Thus, cellulose and its oligosaccharide derivatives have a broad application range, from materials encountered in everyday life, such as paper and textiles, to state-of-the-art specialized materials such as probes for enzyme activity, biosurfactants, nanomaterials and biogels^20–23^.

Due to the important biological roles and application potential of cellulose and its oligosaccharides, the production of cellulose-based structures has attracted much attention. Synthetic chemists are only just starting to report the automated synthesis of β-1,4-linked glucose oligosaccharides, which is challenging due to the required stereo- and regioselectivity^24,25^. The enzymatic synthesis of cellulose *in vitro* is challenging because the natural biosynthetic machinery consists of complex membrane-embedded multi component systems^26^, that are hard to express in functional form. Furthermore, biosynthesis is processive and is biased towards production of very long oligomers^27,28^. Recent advances in employing biocatalysts for *in vitro* cellulose synthesis include production of high molecular weight (DP >200) cellulose or cellulose microfibrils from uridine diphosphate glucose (UDP-Glc) via exploitation of bacterial or plant-derived cellulose synthase (complexes) that are functionally reconstituted in lipid bilayers^29,30^. Native and derivatized cellulose and its oligosaccharides have been generated from glucose-1-phosphate and cellobiose via exploitation of the reversible reaction mechanism of cellodextrin phosphorylases^31–35^, whereby reaction conditions can direct the degree of polymerization of the obtained oligosaccharide mixtures^36,37^. However, irreversible mechanisms of biocatalytic synthesis of soluble cello-oligosaccharides would present an attractive alternative. While progress has been made via glycosynthases, which can generate β-1,4-linked glycosidic linkages using activated Glc-α-F as the donor^38^, it remains challenging to generate natural or derivatized soluble cello-oligosaccharides from natural donors due to a lack of suitable biocatalysts.

Glycosyltransferases synthesize highly regio- and stereospecific glycosidic bonds between glycan acceptors and naturally occurring activated sugar donors with phosphate leaving groups, such as phosphate bound-sugars (e.g. glucose-1-phosphate) or nucleotide-sugars (e.g. UDP-Glc)^39–41^. So far the enzymatic synthesis of Glc-β-1,4-Glc linkages from nucleotide sugars has been restricted to plant and bacterial cellulose synthases and formation of soluble cellulose oligosaccharides by these enzymes has not been reported. However, glycosyltransferases can be promiscuous in acceptor and donor substrates, thereby providing a rich source of biocatalysts with potentially exploitable side reactions. For example, human blood group galactosyltransferase and bovine β4GalT1 have been reported to display activity with multiple UDP-donor substrates including UDP-Glc^42–44^, while *Neisseria meningitidis* LgtB displays activity towards multiple acceptor substrates^45^. The rates of activity on alternative acceptors and donors have in many cases been demonstrated to be sufficient for exploitation of the promiscuous activity in synthesis of glycosides^42,44–48^. In principle, a glycosyltransferase able to accommodate both UDP-Glc as donor and a Glc-terminated structure as acceptor could function as glucose polymerase (Fig. 1), whereby the degree of polymerization can potentially be tuned through optimization of reaction conditions or via enzyme engineering. Substrate promiscuity of glycosyl-transferases is therefore a promising avenue to explore in a rational approach to identify a biocatalyst able to synthesize soluble cellulose oligosaccharides.

**Fig. 1.**
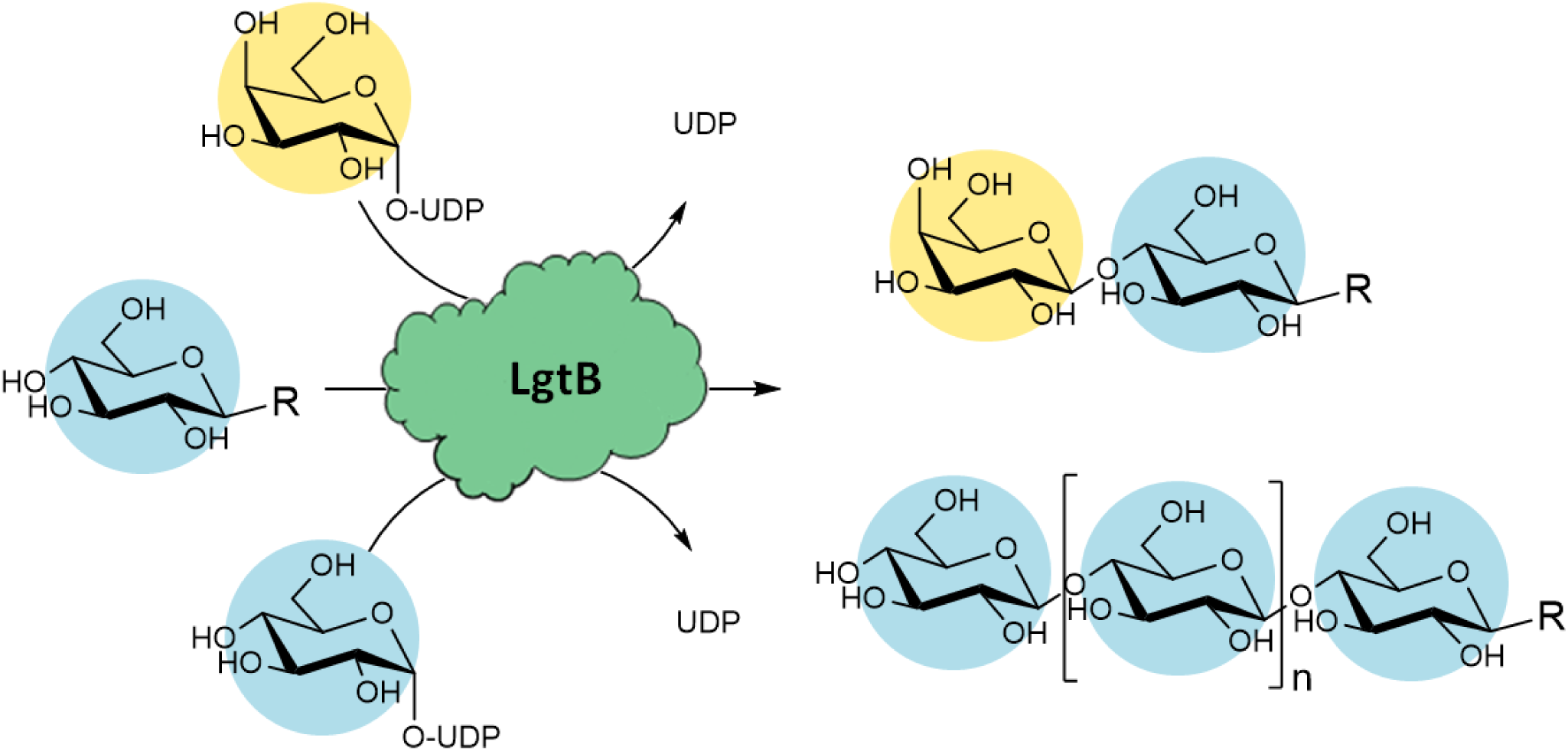
Exploiting promiscuous activity to retask a galactosyltransferase for glucose polymerization. In principle, a glycosyltransferase that accepts UDP-Glc as donor and Glc-terminated structures as acceptor substrate, enables glucose polymerization. Following this rationale, exploration of promiscuous galactosyltransferase activity resulted in identification of a biocatalyst, *Neisseria meningitidis* LgtB, suitable for generation of β-1,4-linked glucose oligosaccharides and their derivatives. This and following diagrams use symbol nomenclature for glycans (SNFG)^52,53^.

Here we describe how the rational exploration of galactosyltransferase substrate promiscuity resulted in the identification of a broad-specificity biocatalyst that functions as a glucose polymerase in vitro. We demonstrate how the enzyme can be applied to the practical synthesis of cellulose-oligosaccharides. Via polymerization onto imidazolium-tagged (ITag-) glycosides^48,49^, we generated cellulose-based probes for enzyme activity that are highly suitable for mass spectrometry-based detection. We demonstrate their utility in substrate-product profiling of hydrolytic and oxidative biomass degrading enzyme activities such as lytic polysaccharide monooxygenases (LPMOs) – enzymes of high importance to industrial biomass conversion for which assays reporting on selectivity are in demand^50,51^. Via incorporation of the polymerase in biocatalytic cascades we gained access to derivatized oligosaccharides with β-1,4-linked glucose motifs, and we demonstrate applicability of these as probes to detect selectivity of cellulolytic enzymes. We anticipate that the polymerase activity of this enzyme will find broad application in biocatalytic strategies to generate glycan structures and probes for glycobiology and wider areas of nanobio-technology.

## Results

### A biocatalyst for glucose polymerization

With the aim to identify a glycosyltransferase capable of generating β-1,4-linked glucose (Glc) oligosaccharides, we assembled a panel of five recombinantly expressed galactosyltransferases and screened it for promiscuous acceptance of Glc-based donor and acceptor substrates. Transfer of UDP-Glc to an acceptor with a terminal Glc-R motif would result in reaction products that can be re-used as an acceptor, thus enabling the desired Glc polymerization. As the acceptor structure can potentially affect the generated glycosidic linkage type^54^, enzymes with a variety of reported specificities were included.

To monitor the activity of the enzyme panel, we employed a sensitive and fast assay based on glycosylation of sugar acceptors labelled with imidazolium-based probes (I-Tags), such as the 4-(1-methyl-3-methyleneimidazolium)benzyl carbamate β-glucoside^48^ ITag-Glc-1 (**1**) (Fig. 2A). Our previous work demonstrated that such I-Tags generate strong signals in mass-spectrometry that dominate the analyte ionization and can be employed for yield estimation in biotransformations^48,55,56^. To assess the scope of the glucose polymerization activity, we chemically synthesized the novel 4-(1-methyl-3-methyleneimidazolium) benzyl β-glucoside ITag-Glc-2 (**2**) (Fig. 2A) in 4 steps (Fig. S5) with a yield of 49%.

**Fig. 2.**
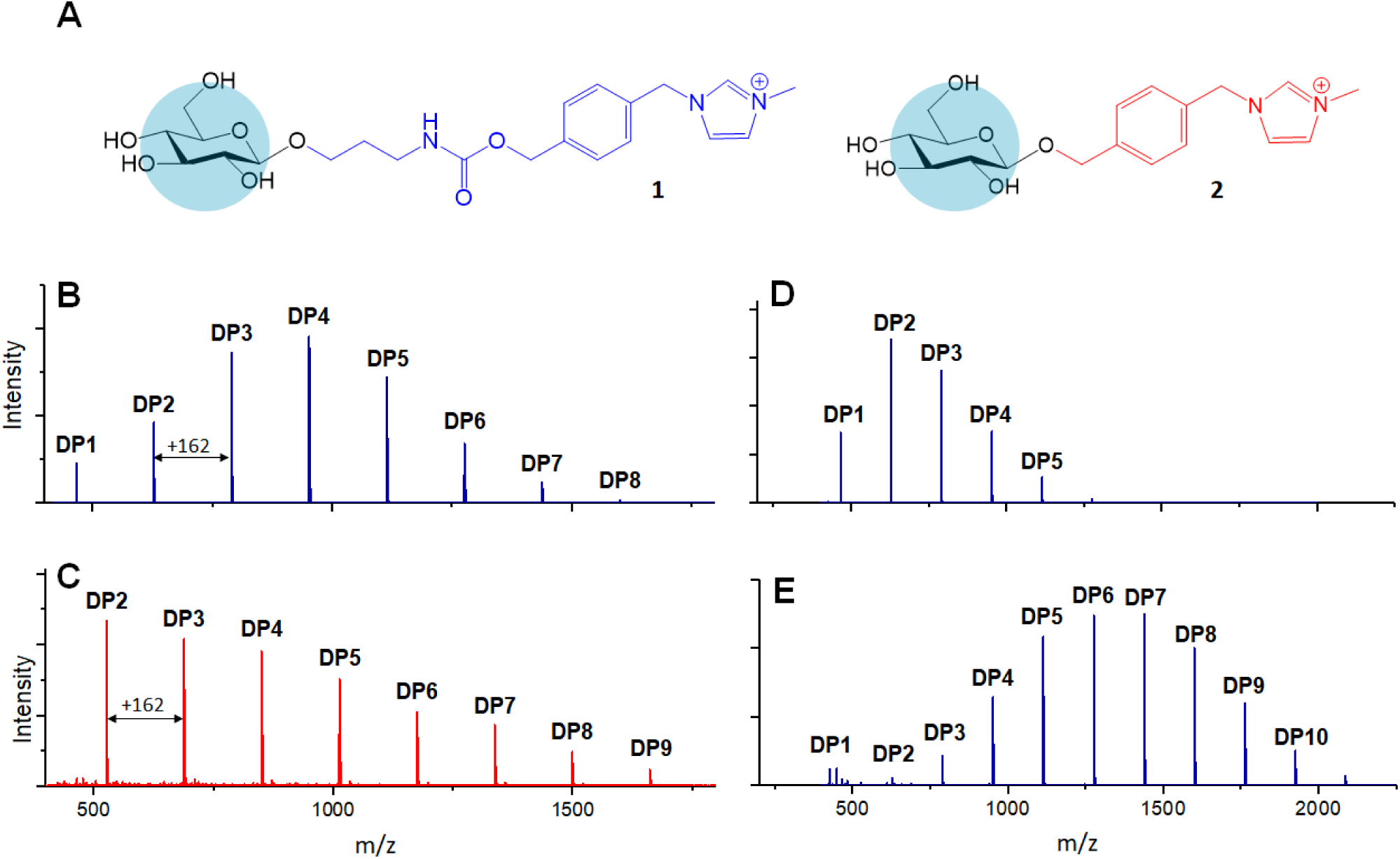
Cello-oligosaccharide envelopes as produced by LgtB via polymerization of glucose onto I-Tagged glucose acceptors. **A**, acceptor substrates for LgtB, glucosides derivatized with I-Tag 1 (**1**) and I-Tag 2 (**2**). **B**, oligosaccharide envelopes as synthesized by LgtB onto acceptor **1** and detected by MALDI-TOF MS. **C**, oligosaccharide envelopes as synthesized by LgtB onto **2** as detected by MALDI-TOF MS. **D, E** length of incubation and UDP-Glc concentration alters length and ratio of oligosaccharides produced, shown after 2 days, 1.5 mM UDP-Glc and 7 days, 15 mM UDP-Glc respectively. Degrees of polymerization are indicated, masses are specified in supplementary information.

The transfer of galactose (Gal) from UDP-Gal to I-Tagged acceptors was detected as an activity displayed by all tested panel members, LgtB, LgtC, LgtH, B4GALT4 and G0PH97 (Fig. S1, S2). In contrast, activity with UDP-Glc was detected only for *Homo sapiens* B4GALT4 and *Neisseria meningitidis* LgtB, with multiple reaction products due to polymerization observed in the latter case. LgtB was therefore identified as the biocatalyst most suitable for our target activity, i.e. glucose polymerization.

After incubation of LgtB with **1** or **2** and an excess of UDP-Glc at 37°C, a range of I-Tagged glucose oligosaccharides with a degree of polymerization (DP) of up to 7 and 9, respectively (Fig. 2B, C) were observed. Adjustment of reaction conditions allowed us to direct the glucose polymerization towards a desired product range (Fig. S6). For example, extending incubation times and increasing donor concentrations resulted in oligosaccharides with higher DPs (Fig. 2D, E). Two-dimensional heteronuclear single quantum coherence NMR (2D HSQC NMR) analysis of purified I-Tag Glc oligosaccharides generated from (**1**) confirmed that the Glc residues were connected by β-1,4-glycosidic linkages (Fig. S7), demonstrating the strict selectivity in anomeric configuration and position of the glycoside linkage that is formed by LgtB.

The glucose polymerization reaction was not restricted to the ITag-Glc acceptors; LgtB was able to polymerise Glc onto a broad range of acceptor substrates (Table S1) including native cello-oligosaccharides and derivatives with reducing end conjugates such as Glc_(n)_-pNP (Fig. S4). This is in line with the reported broad acceptor substrate scope of this enzyme, which has been exploited for chemo-enzymatic synthesis of β-1,4-linked galactosides incorporating e.g GlcNAc(-pNP), Man-pNP, Glc(-pNP) and various C2-derivatives^45–48^. No activity was found using Gal, xylose (Xyl), arabinose (Ara), lactose (Gal-β-1,4-Glc) and trehalose (Glc-α-1,1-Glc) as acceptors. Acceptor substrates thus require an equatorial configuration of the C4 -OH group and the presence of a C6 -OH group while both the C1 and C2 substitutions are highly flexible. Further examining the donor scope of LgtB, transfer of Xyl and GalNAc but not GlcNAc from their respective UDP-conjugates to acceptors was also detected, although with lower yields (Fig. S3).

The discovery of the extended biosynthetic scope of LgtB opens up the possibility of including cello-oligosaccharide biosynthesis in biocatalytic cascades, such as those enabling regeneration of UDP-glucose. We demonstrated this principle by combining LgtB in a one pot, two enzyme cascade (Fig. 3) with *Solanum lycopersicum* sucrose synthase (SLSUS6), hereafter SuSy. In the presence of UDP, SuSy converts relatively inexpensive sucrose into fructose and UDP-Glc^57^, this biocatalyst is therefore widely employed for the synthesis of nucleotide sugars and glucosides^58,59^. Combining SuSy in incubations with LgtB and acceptor substrates **1** or **2** resulted in the formation of I-Tagged cello-oligosaccharides (Fig. S8) with DP ranges and ratios similar to those observed in LgtB reactions using UDP-Glc directly as donor substrate. These results demonstrate how UDP-Glc generated via SuSy activity from sucrose, can be used as a donor substrate by LgtB, resulting in a successful integration of both enzymes in a one-pot biocatalytic cascade that converts sucrose to I-Tagged cello-oligosaccharides, and points to the polymerization of glucose from cheap sucrose starting materials.

**Fig. 3.**
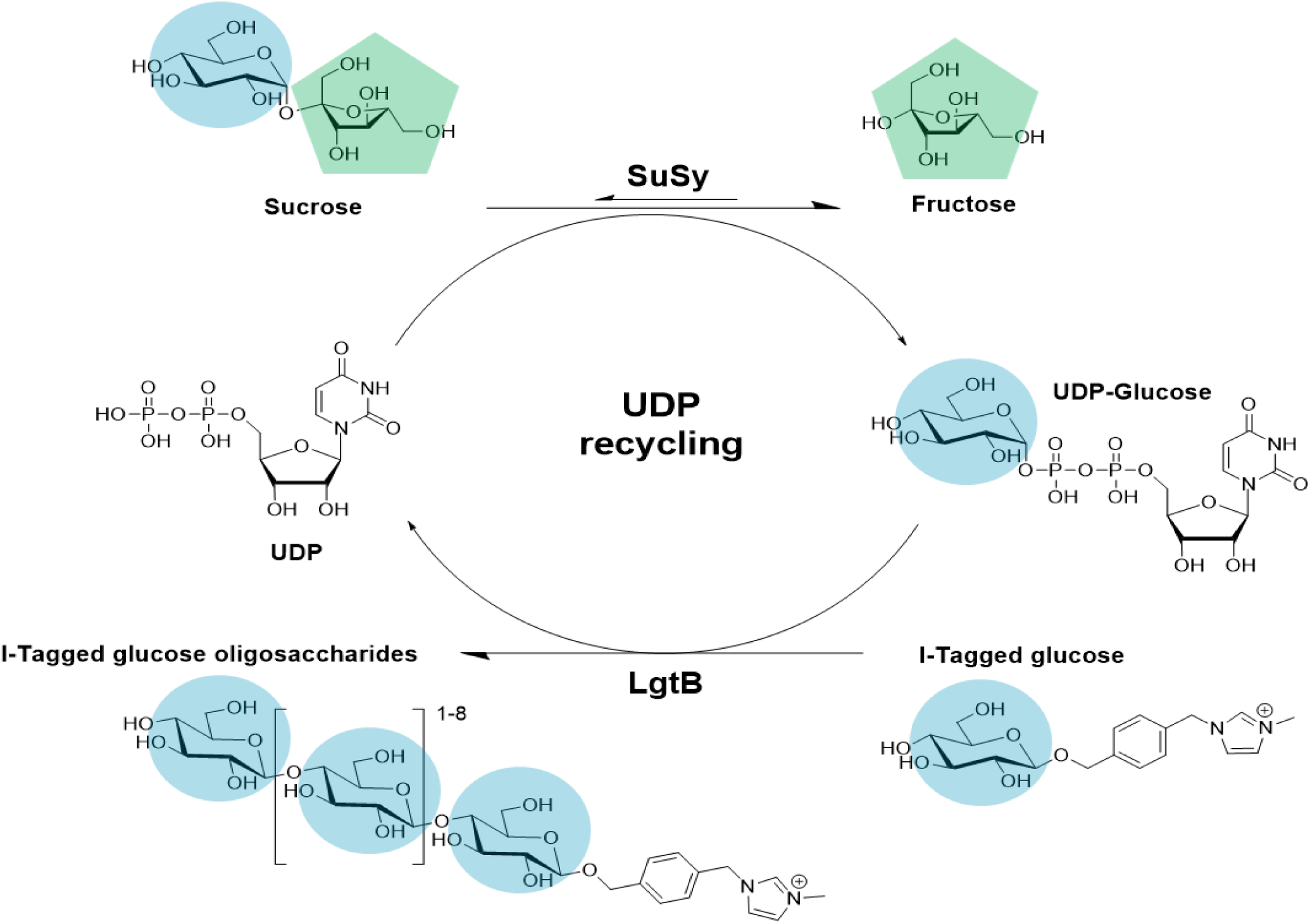
Biocatalytic cascade enabling effective coupling of UDP-glucose regeneration with glucose polymerization. LgtB and sucrose synthase (SuSy) were utilised in a one-pot reaction for the production of I-Tagged cello-oligosaccharides.

Taken together, LgtB was identified as a promiscuous biocatalyst with glucose polymerase activity that can be exploited for facile polymerization of glucose onto a broad range of acceptors, resulting in native and derivatized cello-oligosaccharides that are highly relevant in the development of materials or probes for cellulolytic enzymes.

### Exploiting LgtB to synthesize probes for LPMO activity profiling

We exploited I-Tagged glucose oligosaccharides, generated via LgtB activity, to rapidly profile the catalytic activity and substrate preferences of lytic polysaccharide monooxygenases (LPMOs), a family of copper dependent enzymes that oxidatively degrade recalcitrant polysaccharides such as cellulose, chitin and starch^60–63^. LPMOs utilize molecular oxygen or hydrogen peroxide to insert a single oxygen atom into the C1-H and/ or C4-H bond of saccharide substrates^61,64,65^ generating lactone or 4-ketoaldose products that can spontaneously convert to aldonic acids or 4-gemdiol-aldoses, respectively (Fig. 4A). LPMOs have attracted considerable commercial interest as auxiliary enzymes for the cost-effective production of biofuels from lignocellulosic feedstocks, and also hold great promise for the selective functionalization of carbohydrates for the development of (ligno)cellulose-based chemicals and materials. As such, there is great interest in developing sensitive high-throughput assays that report on LPMO selectivity for C1-vs C4-oxidation, and allow identification of enzymes with activity on soluble low- to medium-molecular weight oligomers. This can in principle be achieved by employing I-Tagged probes in LPMO activity assays, followed by mass spectrometry-based detection of reaction products. The cationic nature of I-Tagged probes avoids formation of sodium or potassium adducts^48^ that have been demonstrated to complicate mass spectrum interpretation in LPMO activity assays^66^.

**Fig. 4.**
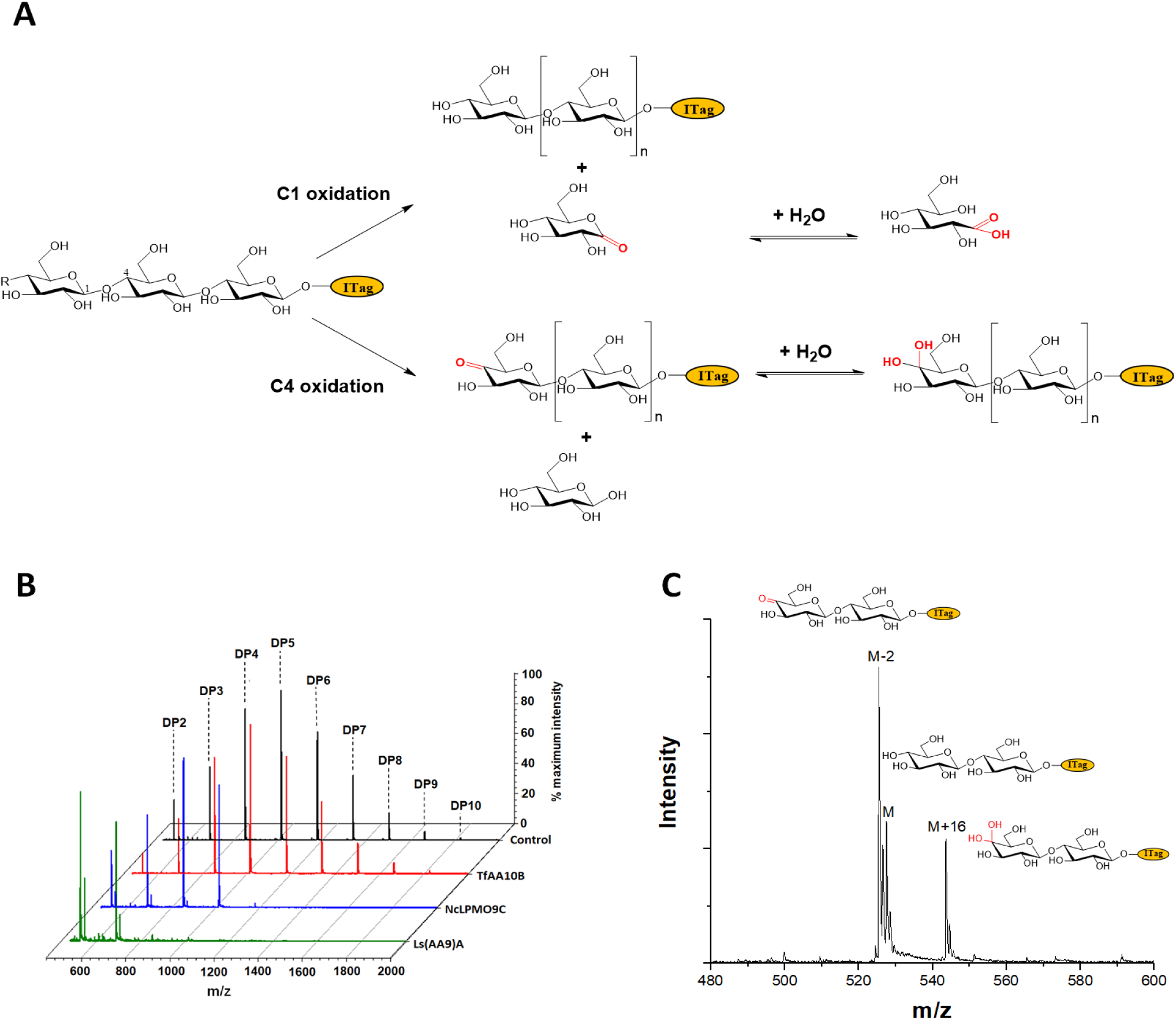
I-Tagged cello-oligosaccharides as informative probes for LPMO activity. **A**, the C1- and C4-oxidative activity of LPMO on I-Tagged oligosaccharides results in distinctive reaction products that can be easily identified by MALDI-TOF MS. **B**, diagram demonstrating selective activity on I-Tagged glucose oligosaccharide substrates by a panel of LPMOs. **C**, the products of *Nc*LPMO9C activity on I-Tagged cello-oligosaccharides include those with a mass corresponding to C4-ketone (M-2) and gemdiol (M+16) disaccharides, confirming C4-oxidative activity.

A small panel of LPMOs was produced in *E. coli* as the expression host; *Aspergillus oryzae* LPMO of CAZy family AA11 A*o*(AA11)^60^, *Lentinus similis Ls*(AA9)A^61,65^, *Neurospora crassa Nc*LPMO9C^51,62,67^ and *Thermobifida fusca Tf(*AA10)B^68^ (see SI for detailed protocols and accession numbers). Biotransformations were performed with purified enzymes, using an envelope of I-Tagged glucose oligosaccharides (DP 1-10) in the presence of H_2_O_2_ and ascorbate as a reducing agent. The distribution of labelled oligosaccharides was unchanged following incubation with either *Tf(*AA10)B or *Ao*(AA11) (Fig. 4B, S9), demonstrating that these enzymes are not active towards the oxidation of soluble cello-oligosaccharides with DP < 10. In contrast, reactions with *Nc*(AA9)C and *Ls*(AA9)A led to complete consumption of oligosaccharides with DP ≥ 6 and 4, respectively (Fig. 4B), with concomitant formation of labelled C4-oxidized products (DP 2-4 and DP 2-3, respectively) with molecular weights of [M-2] (C4-ketone) and [M+16] (C4-gemdiol) (where M is the molecular weight of the corresponding non-oxidized labelled carbohydrate) (Fig. 4C). These combined data are consistent with the previously reported substrate profiles of *Tf(*AA10)B, *Ao*(AA11), *Nc*(AA9) C and *Ls*(AA9)A, and highlight the utility of I-Tagged glucose oligosaccharides envelopes for rapidly profiling LPMO activity.

### Facile chemo-enzymatic synthesis of probe derivatives and their application

To further explore the application of I-Tagged cello-oligosaccharides as enzyme activity probes, we initially verified activity of a commercially obtained β-glucosidase from Almond and of a mixture of cellulases from *Aspergillus niger* on the probes (Fig. 5A-C). The β-glucosidase has been reported to cleave glucose residues from di- and oligosaccharides in an exo-acting fashion whereas the cellulases function mainly in a endo-acting manner^69^. After 1 h incubation with β-glucosidase at 37 ^°^C, analysis of reaction products via MALDI-TOF MS resulted in detection of only a single peak of *m/z* 466, indicating that as expected the enzyme hydrolyzed high-DP oligosaccharides, leaving only a single Glc residue attached to the ITag (Fig. 5C). A 2 h incubation with the cellulases resulted in a similar product (Fig. 5B), whereas no changes were detected after incubation with a negative control containing only buffer. These results confirm that the enzymatically synthetized I-Tagged cello-oligosaccharides are suitable probes to detect glycoside hydrolase activity.

**Fig. 5.**
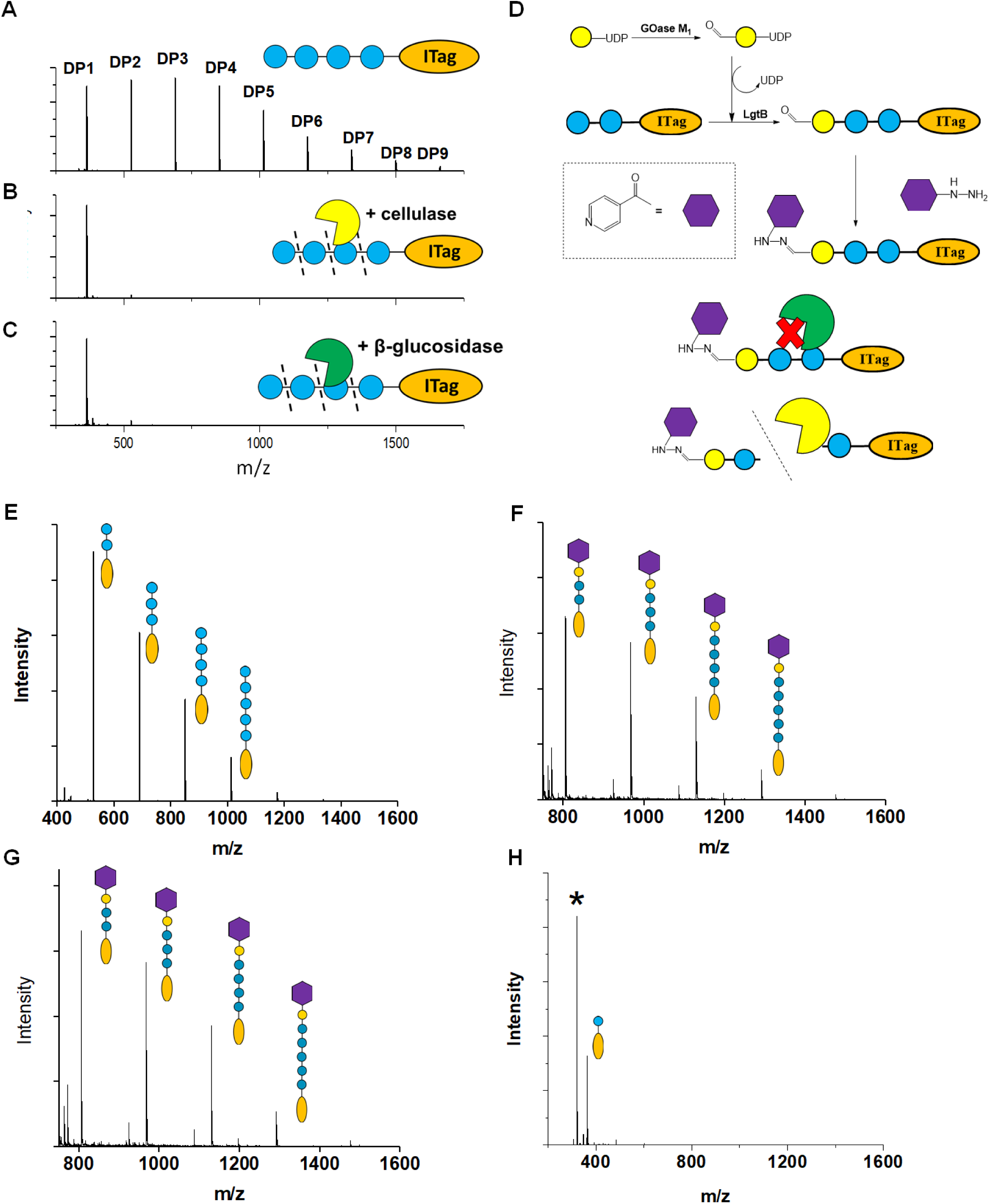
Chemo-enzymatic derivatization of cello-oligosaccharides to selective endo-cellulase substrates. **A**, Representative MALDI-TOF MS spectra demonstrating I-Tagged cello-oligosaccharides after incubation without any cellulolytic enzyme **B**, with *A. niger* cellulase and **C**, with β-glucosidase from almond. **D**, derivatization strategy employed to protect LgtB product termini against hydrolytic activity; **E**, MALDI-TOF MS spectra of the envelope of purified I-Tagged cello-oligosaccharides; **F**, probe envelope obtained after ligation to oxidised galactose and nicotinic hydrazide; **G**, probe envelope after incubation with β-glucosidase and β-galactosidase; **H**, probe envelope after incubation with *A. niger* cellulases. Asterix indicates a product corresponding to a glycosylated hydrazide.

Selective detection of endo-cellulase activity is in principle enabled by generating an enzyme substrate that cannot be hydrolysed via hydrolysis of its termini. Following this principle, progress has been made in generating ‘blocked’ substrates for endo-cellulases and endo-xylanases that are compatible with colorimetric assays (e.g detection of released 2-chloro-4-ni-trophenol)^70,71^. Here we expand the range of ‘blocked’ substrates to probes that are compatible with fast and sensitive detection of activity via MALDI-TOF MS. We demonstrate how LgtB can effectively be incorporated in chemo-enzymatic strategies to derivatize or tailor cellulose structures, opening up access to chemical building blocks or tailored enzyme probes.

‘Blocked’ I-Tagged glucose oligosaccharides were generated via a chemo-enzymatic approach based on creation of a bio-orthogonal aldehyde on the non-reducing termini of oligosaccharides, which is subsequently derivatized with a nicotinic hydrazide group (Fig. 5D). Enzymatic oxidation of UDP-Gal to UDP-Gal_ox_ was achieved using *F. graminearium* galactose oxidase (GOase) variant M_1_ ^72^, reaching full conversion after overnight incubation at 25 °C (Fig. S10). I-Tagged oligosaccharides were generated with LgtB, purified (Fig. 5E), and the UDP-Gal_ox_ was subsequently transferred onto their non-reducing termini via overnight incubation with LgtB. The introduction of this bio-orthogonal aldehyde enables selective derivatization of the terminal Gal moieties via previously described methodologies^73,74^. Here, nicotinic hydrazide was ligated with the C6-Gal aldehyde to generate ‘blocked’ oligosaccharides (Fig. 5F). To demonstrate the necessity of the I-Tag at the reducing end, the same derivatization strategy was applied to cellohexaose. Incubation of the Gal_ox_ containing cellohexaose with nicotinic hydrazide generated a double ligated species with the nicotinic hydrazide ligating at the aldehyde of the Gal_ox_ as well as the aldehyde at the free reducing end. This generated a particularly complex mixture, with which it would prove difficult to assess the effectiveness of hydrolases (Fig. S11).

With the blocked substrates in hand, we confirmed protection of the substrates against exo-acting enzyme activity via incubations of this envelope of probes and intermediate products with exo-acting enzymes. The addition of a terminal galactose was sufficient to protect the cello-oligosaccharides oligosaccharides against β-glucosidase activity, while the derivatization with the hydrazide group was required and sufficient to protect structures against β-galactosidases from a range of glycoside hydrolase families (GH1, GH42 and GH50) (Fig. 5G, Fig. S12). Finally, we employed the blocked substrates to monitor endo-acting cellulases for activity, and validated the presence of cellulolytic activity by MALDI-TOF MS (Fig. 5H). The endo-cellulase reaction product profile was similar to those of the same enzymes incubated with I-Tagged oligosaccharides with a free non-reducing end. Taken together, we have developed a probe for the selective detection and profiling of endo-cellulase activity via MALDI-TOF MS, demonstrating how LgtB mediated synthesis of cello-oligosaccharides can be successfully incorporated into chemo-enzymatic derivatization strategies.

## Conclusion

We have demonstrated that glycosyltransferase LgtB has a broad substrate scope that includes polymerase activity using UDP-Glc as donor and Glc-terminated glycosides as acceptors. This promiscuous activity can be exploited to polymerize glucose via β-1,4-glycosidic linkages. Synthesis of such cello-oligosaccharides onto acceptors derivatized on the reducing termini with a variety of chemical handles resulted in functionalized oligosaccharides. As the LgtB enzyme can be obtained via facile heterologous expression in *E. coli*, these findings enable straightforward synthesis of enzyme probes and generate new strategies for (chemo)enzymatic synthesis of cello-oligosaccharide derivatives. As proof-of-principle we exploited LgtB to generate I-Tagged cello-oligosaccharides, which were employed in a sensitive and robust MALDI-TOF MS-based assay to report on substrate use of cellulolytic enzymes. In addition to detecting enzyme activity and enabling profiling of substrate specificity, the probes allow facile and reliable discrimination between products from LPMO’s C1 and C4 oxidation reactions. The resulting semi-high throughput profiling of LPMO activity in combination with the facile bacterial LPMO expression system opens up the exciting possibility of LPMO enzyme engineering. The incorporation of LgtB catalytic activity in a broader chemo-enzymatic strategy to derivatize or tailor cello-oligosaccharides has the potential to open up access to chemical building blocks or tailored enzyme probes. We demonstrated this via the generation of probes to selectively detect endo-acting cellulolytic enzyme activities, achieved via chemo-enzymatic derivatization of the non-reducing cello-oligosaccharide termini to protect them against hydrolysis.

## Acknowledgements

This study was funded by the BBSRC, EPSRC and InnovateUK: IBCatalyst programme via BB/P011462/1 (JvM), DTP EP/N509565/1 (JvM), BB/M027023/1 (APG), BB/M029034/1 (SLF), BB/L013762/1 (SLF), BB/M028836/1 (SLF), BB/M027791/1 (SLF), the Marie Sklodowska-Curie Innovative Training Network (H2020-MSCA-ITN-2014-ETN-642870), European Union (CarboMet 737395), and the European Research Council (ERC) Starter Grant 757991 (APG), CoG 648239 (MCG), BB/M028976/1 (MCG), 788231-Pro-grES-ERC-2017-ADG (SLF). We are grateful for the help provided by Dr Matthew Cliff for NMR analysis.

