## supporting information for "In vitro biosynthesis of poly-β-1,4-glucan derivatives using a pro-miscuous glycosyltransferase"

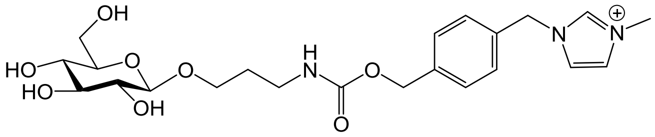

Molecular Weight: 466.51

**1**

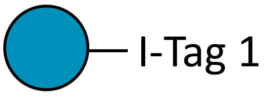

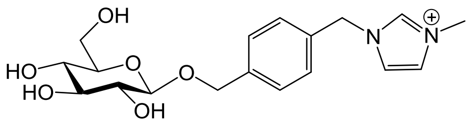

Molecular Weight: 365.41

**2**

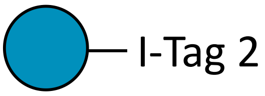

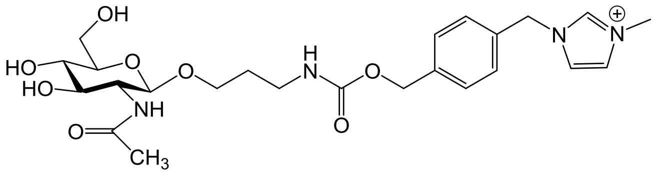

Molecular Weight: 507.56

**3**

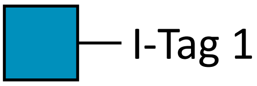

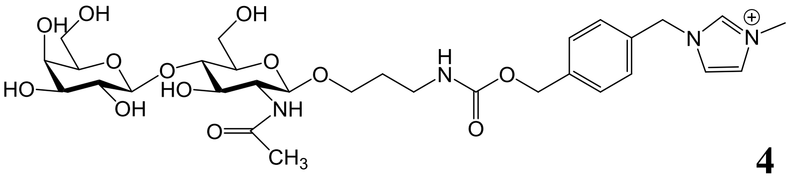

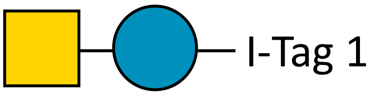

**4**

Molecular Weight: 669.70

**Supplementary figure 1 | I-Tagged oligosaccharides used in this study**

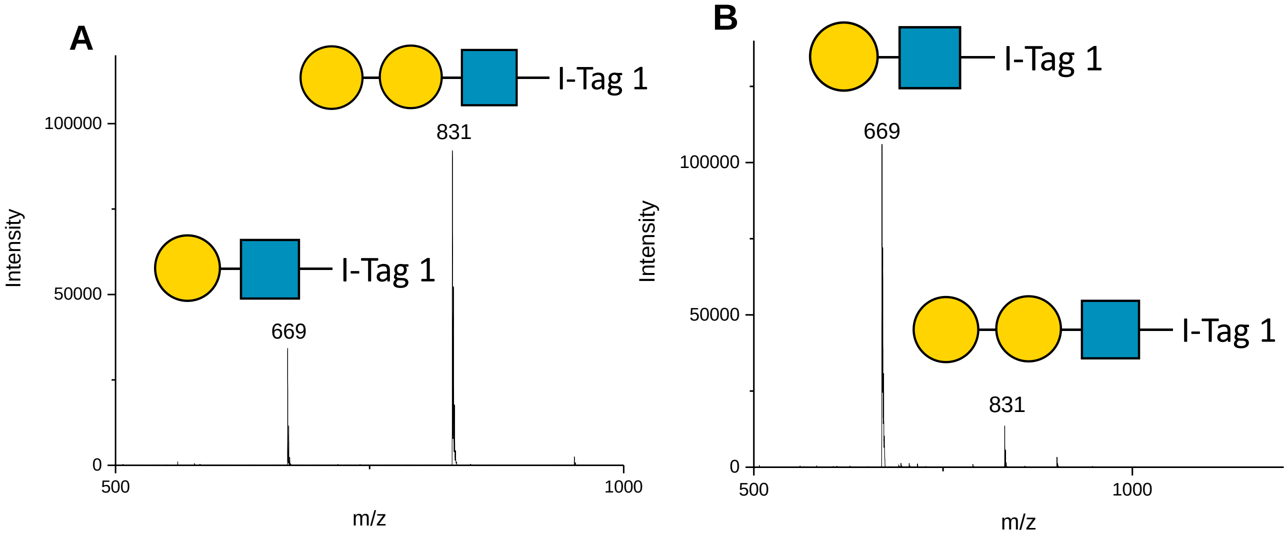

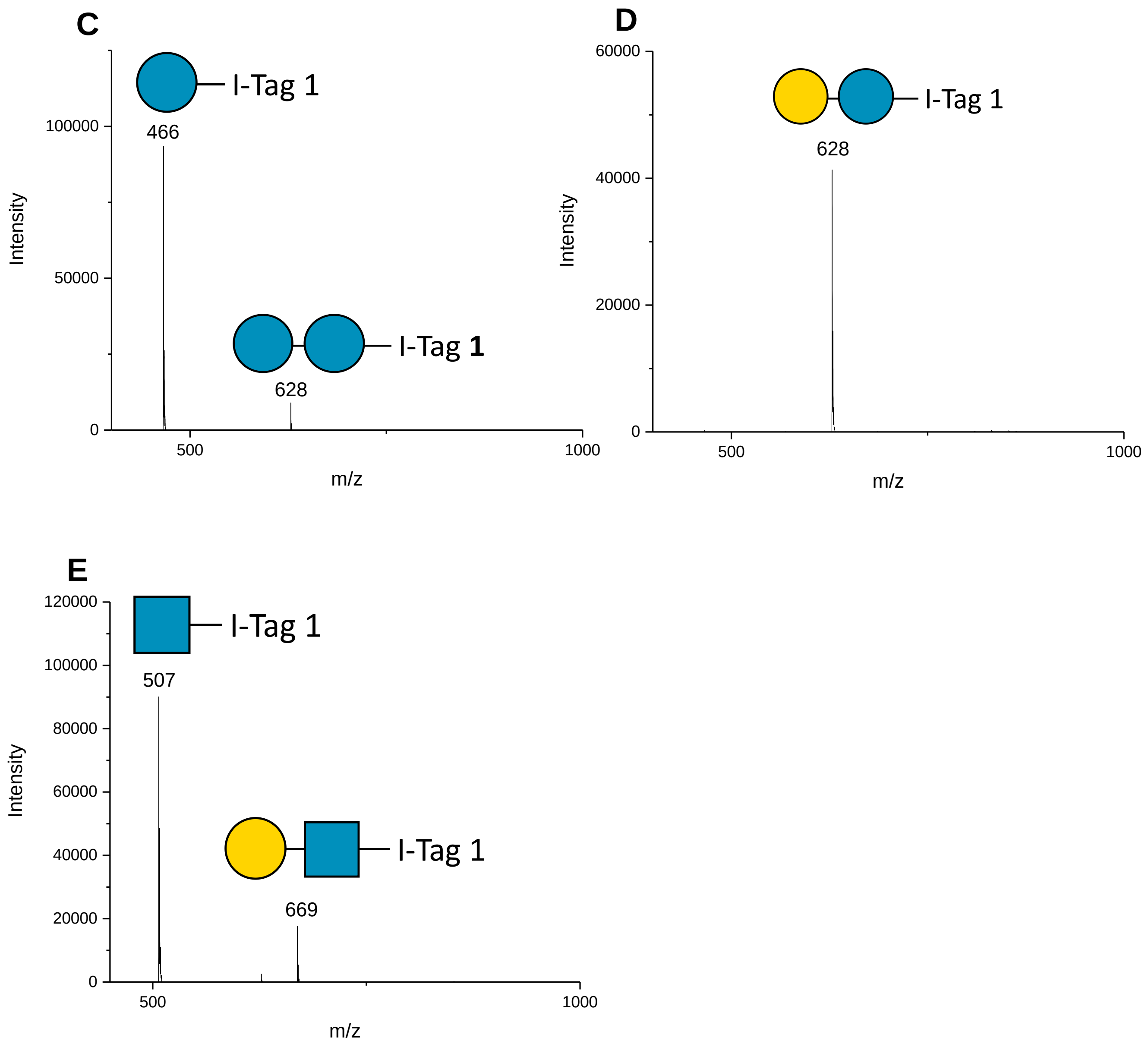

**Supplementary figure 2| Transgalactosylation of I-Tagged glycosides during activity screening.** Transgalactosylation activity of LgtC (**A**) and G0PH97 (**B**) against LacNAc-ITag **4**. A peak of *m/z* 831 was observed for both, corresponding to galactosylated LacNAc-ITag. B4GALT4 demonstrated both UDP-Glc (**C**) and UDP-Gal (**D**) transfer onto Glc-ITag. LgtH (**E**) demonstrated transgalactosylation activity against GlcNAc-ITag **3**. A peak of *m/z* 669 was observed, corresponding to galactosylated GlcNAc-ITag (LacNAc-ITag).

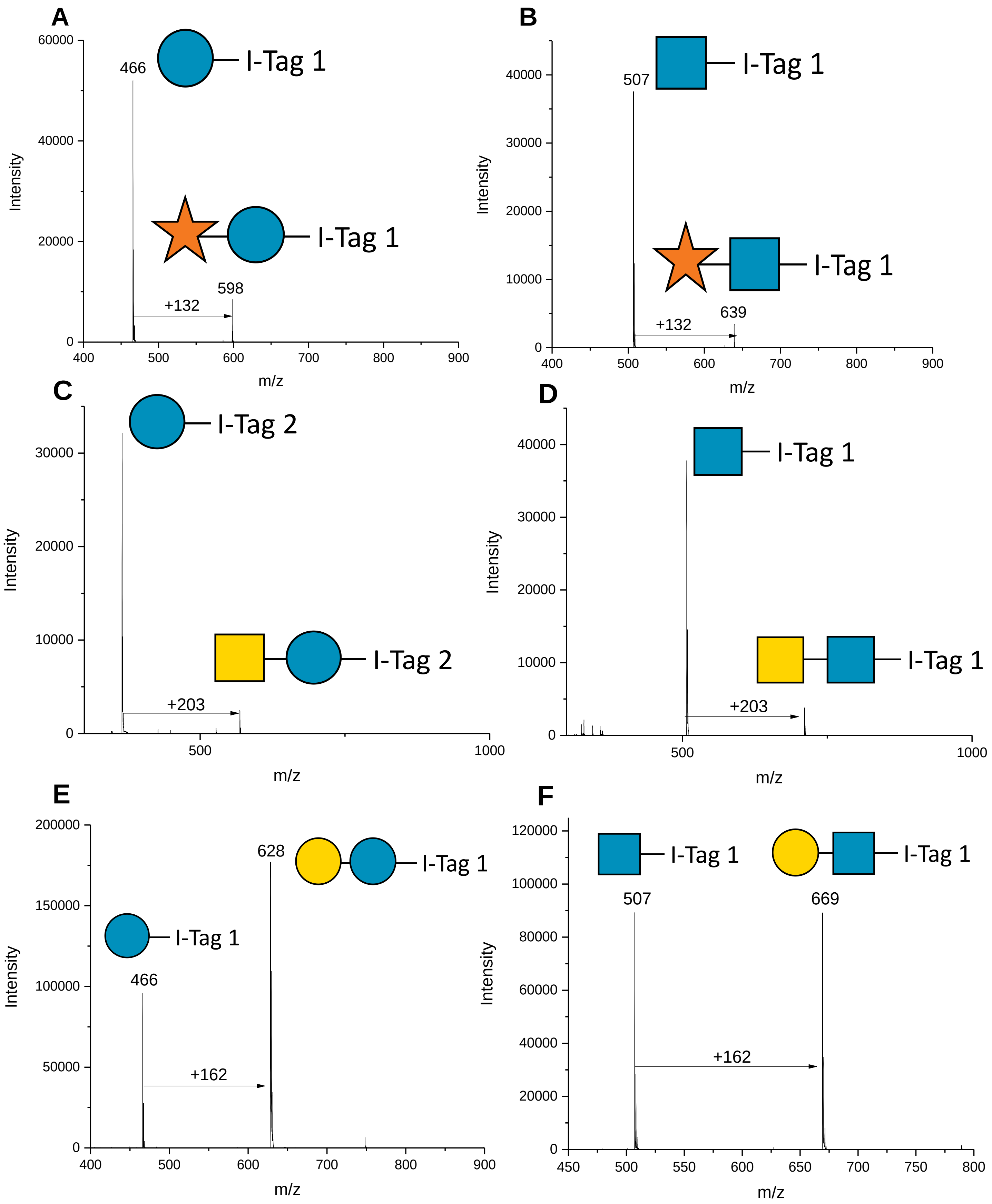

**Supplementary figure 3 | Donor scope assessment of LgtB with I-Tagged acceptors**

LgtB was able to transfer sugar from their respective UDP-conjugate when incubated with (**A** and **B**) UDP-Xyl; (**C** and **D**) UDP-GalNAc and (**E** and **F**) UDP-Gal onto I-Tag-Glc and I-Tag-GlcNac, respectively.

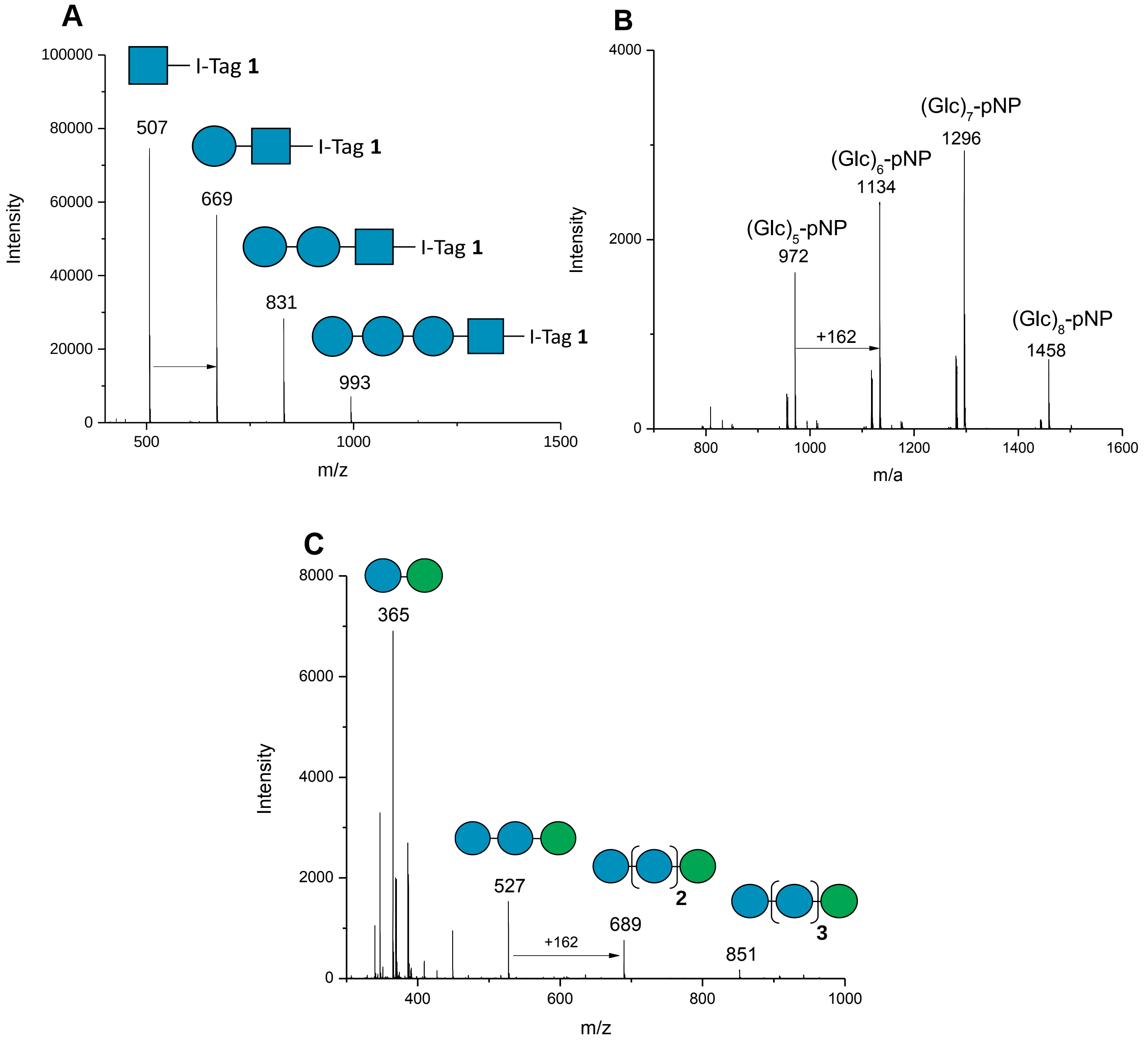

**Supplementary figure 4 | Glucose polymerization occurs onto a range of acceptors**
Glc polymerisation by LgtB was observed via MALDI-TOF MS when using as acceptor substrates **A,** ITag-GlcNAc, **B**, 4-Nitrophenyl β-D-glucose (Glc-pNP) and **C**, Mannose. Products are detected as [M] (for **A**) or [M+Na$^+$] (for **B**, **C**)

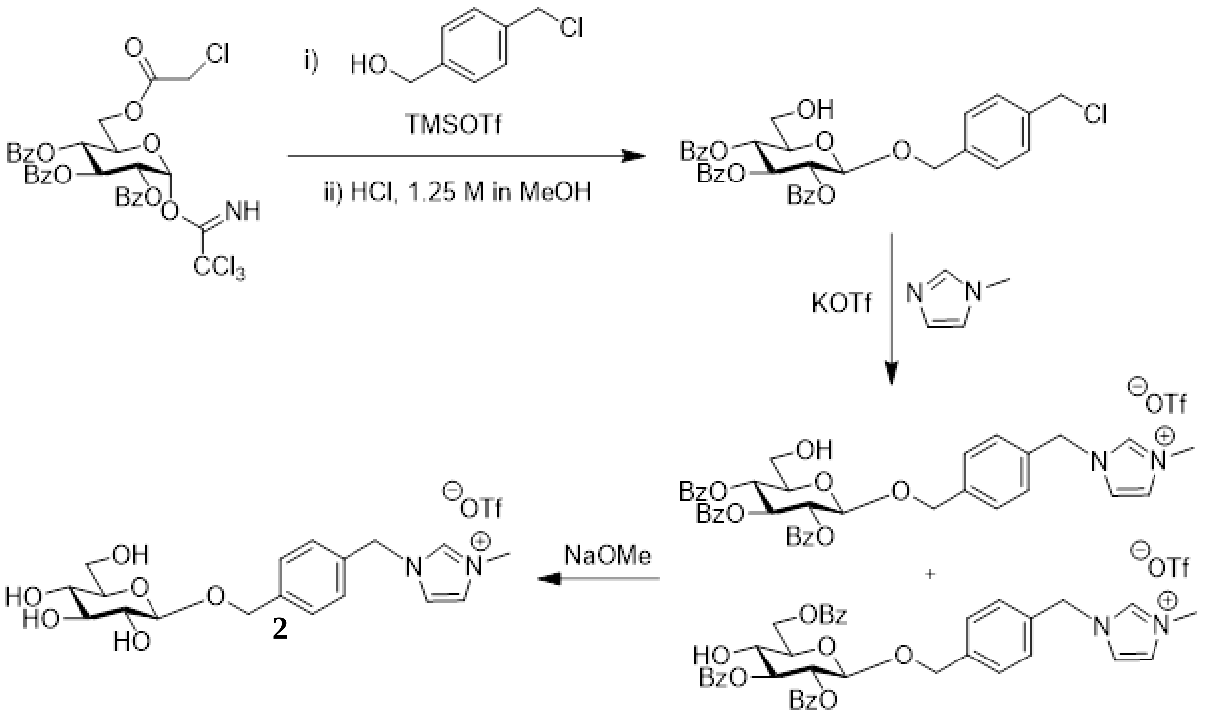

**Supplementary figure 5 | Synthesis of 4-(1-Methyl-3-methyleneimidazolium)benzyl β-D-glucopyranoside trifluoromethanesulfonate (2)**

Compound 2 was synthesized from 4-(chloromethyl)benzyl alcohol and glycosyl donor 2,3,4-tri-O-benzoyl-6-O-chloroacetyl-α-D-glucopyranosyl trichloroacetimidate in 4 steps with a yield of 49%, as described in detail in the methods section. Products were identified as follows: **IR** $\nu_{max}$/cm$^{-1}$ 3419br (OH), 3156w, 3113w, 2968w, 2929w, 1577, 1452, 1416, 1253s, 1225, 1160, 1076, 1028s, 758, 638, 574, 517;  - 24° [*c* 1.08, MeOH]. **$^{1}$H NMR** $\delta_H$ (500 MHz, Methanol-$d_4$) 8.95 (1 H, s, NC*H*N), 7.58 (1 H, d, *J* 2.0, NC*H*CHN), 7.56 (1 H, d, *J* 2.0, NCHC*H*N), 7.53 – 7.48 (2 H, m, $H_{arom}$), 7.43 – 7.38 (2 H, m, $H_{arom}$), 5.39 (2 H, s, NC$H_2$), 4.94 (1 H, d, *J* 12.3, (C-1)OC*H*H), 4.70 (1 H, d, *J* 12.3, (C-1)OCH*H*), 4.35 (1 H, dd, *J* 7.7, 0.9, H-1), 3.92 (3 H, s, NC$H_3$), 3.89 (1 H, dd, *J* 12.0, 2.1, H-6a), 3.68 (1 H, dd, *J* 11.8, 5.5, H-6b), 3.37 – 3.23 (4 H, m, H-2, H-3, H-4, H-5); **$^{13}$C NMR** $\delta_C$ (126 MHz, Methanol-$d_4$) 140.64 (4° $C_{arom}$ (CH$_2$O(C-1))) , 134.44 (4° $C_{arom}$ (CH$_2$N)), 129.91, 129.65 ($C_{arom}$), 125.21 (NCH*C*HN), 123.60 (N*C*HCHN), 103.41 (C-1), 78.10, 78.04, 75.11, 71.66 (C-2, C-3, C-4, C-5), 71.07 ((C-1)O*C*H$_2$), 62.77 (C-6), 53.83 (N*C*H$_2$), 36.52 (N*C*H$_3$); **m/z** (ESI-HRMS) $C_{18}H_{25}N_2O_6^+$ ([M – OTf]$^+$) calculated: 365.1707; found 365.1712; (TLC-MS- (ESI)) $CF_3O_3S^-$ ([OTf]$^-$) calculated 149.0; found 148.8.

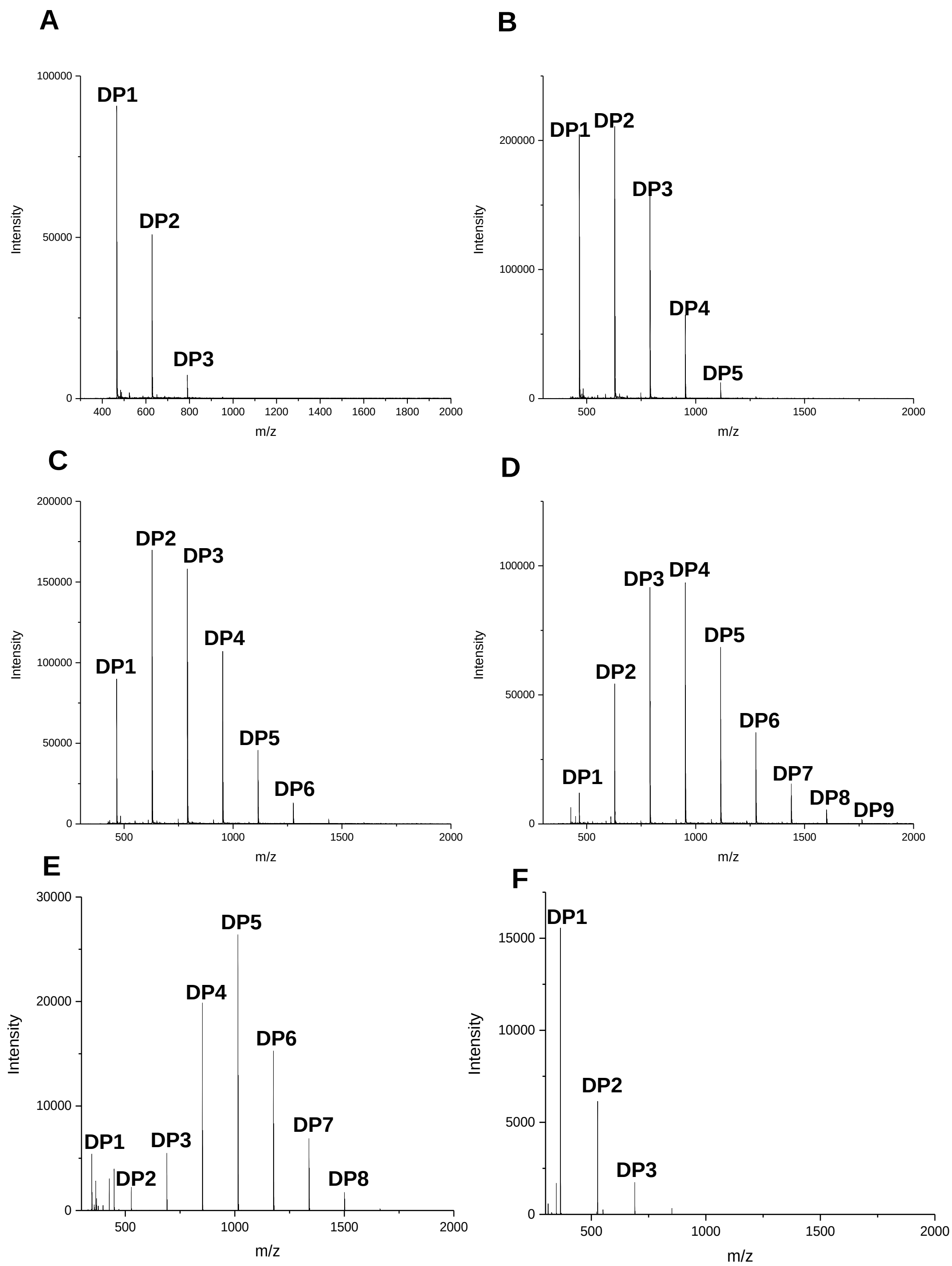

**Supplementary figure 6 | Modification of reaction conditions drives oligosaccharide profile distribution**

Product distribution broadens and degree of polymerization increases during polymerisation of Glc onto **1** after 96 h in response to LgtB concentration with **A** 0.17 mg $ml^{-1}$ LgtB **B** 0.35 mg $ml^{-1}$ LgtB, **C** 0.85 mg $ml^{-1}$ LgtB, **D** 1.7 mg $ml^{-1}$ LgtB. Starting concentration of acceptor affects oligosaccharide distribution, reactions analyzed after 24 h with 1.7 mg $ml^{-1}$ LgtB and **E** 0.2 mM **2** and **F** 0.5 mM **2**.

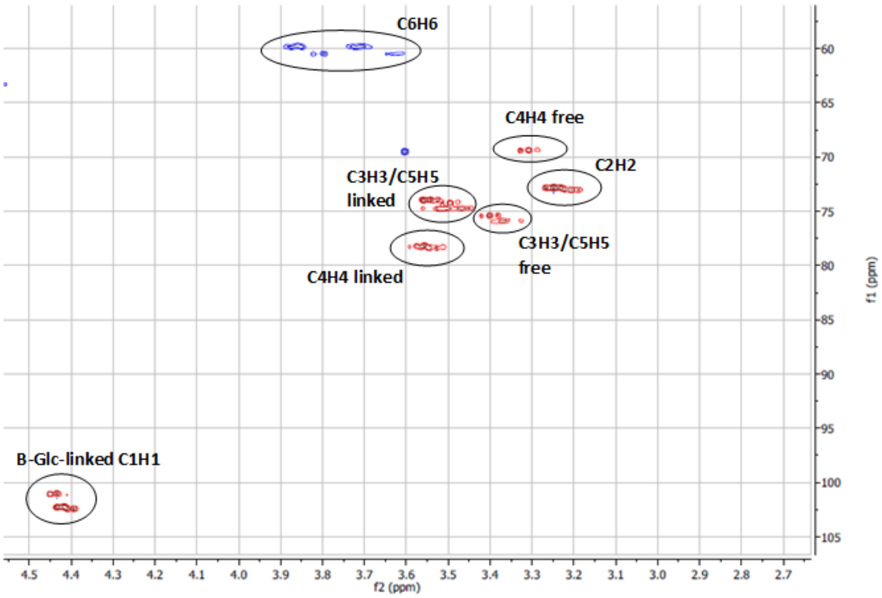

**Supplementary figure 7 | HSQC confirmed that I-Tagged oligosaccharides generated by LgtB consist of β1,4-linked glucose**

$^{1}H$-$^{13}C$ gradient-selected sensitivity-enhanced multiplicity-edited HSQC of purified I-Tagged oligosaccharides generated by LgtB demonstrated the presence of β1,4-linked glucose.

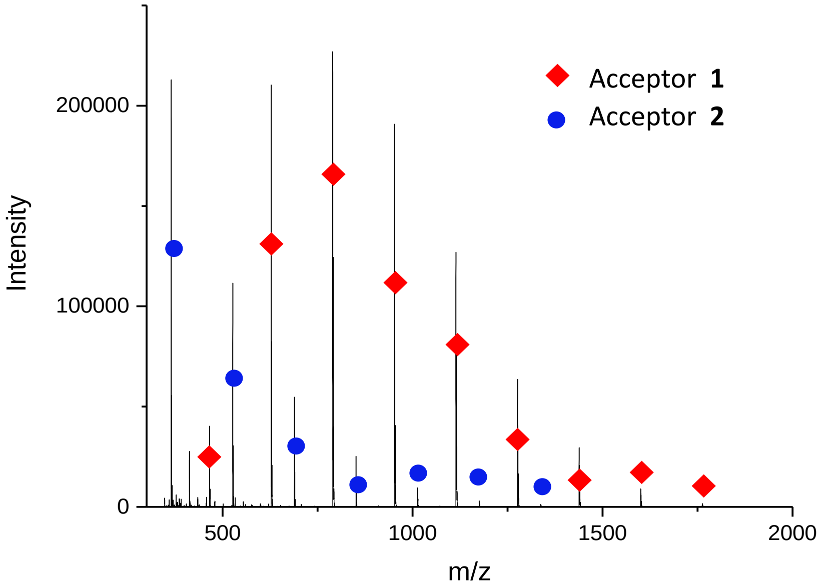

**Supplementary figure 8 | SuSy cascade enables recycling of UDP for donor production**

Superimposed MALDI-TOF spectra of oligosaccharide formation using the SuSy UDP recycling system, with **1** and **2** as acceptors. Degree of polymerisation masses for **1** (red) as follows DP 1 (466), DP2 (628), DP3 (790), DP4 (952), DP5 (1114), DP6 (1276), DP7 (1438), DP8 (1600), DP9 (1762). Degree of polymerisation masses for **2** (blue) as follows DP 1 (365), DP2 (527), DP3 (689), DP4 (851), DP5 (1013), DP6 (1175), DP7 (1337).

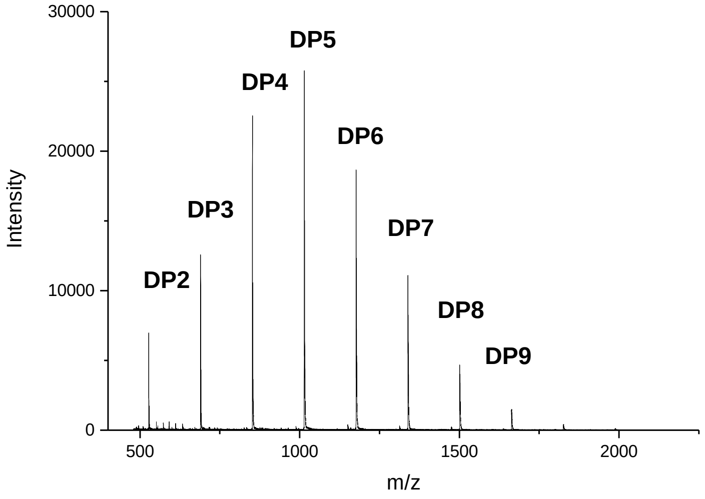

**Supplementary figure 9 | *Ao*(AA11) LPMO digestion of I-Tagged glucose oligosaccharides**

MALDI-TOF MS profile after I-tagged glucose oligosaccharide mixture incubated with LPMO *Ao*(AA11), as expected from previous studies[60] no activity was detected.

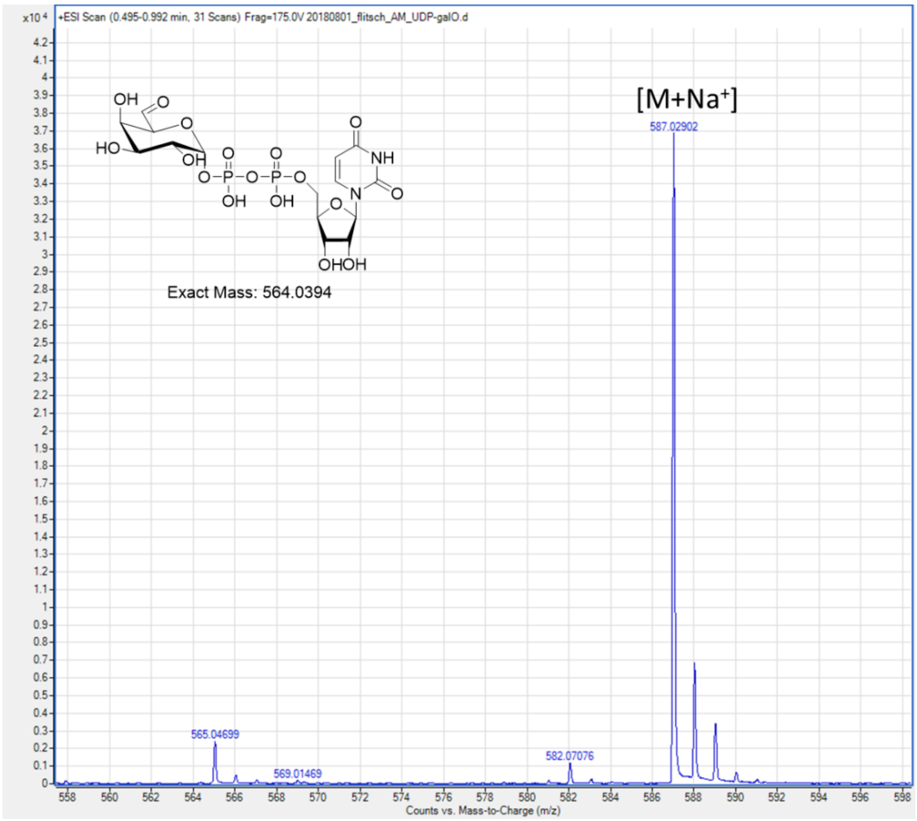

**Supplementary figure 10 | Production of oxidized UDP-Gal**
HRMS data showing production of oxidized UDP-Gal via $M_1$ GOase.

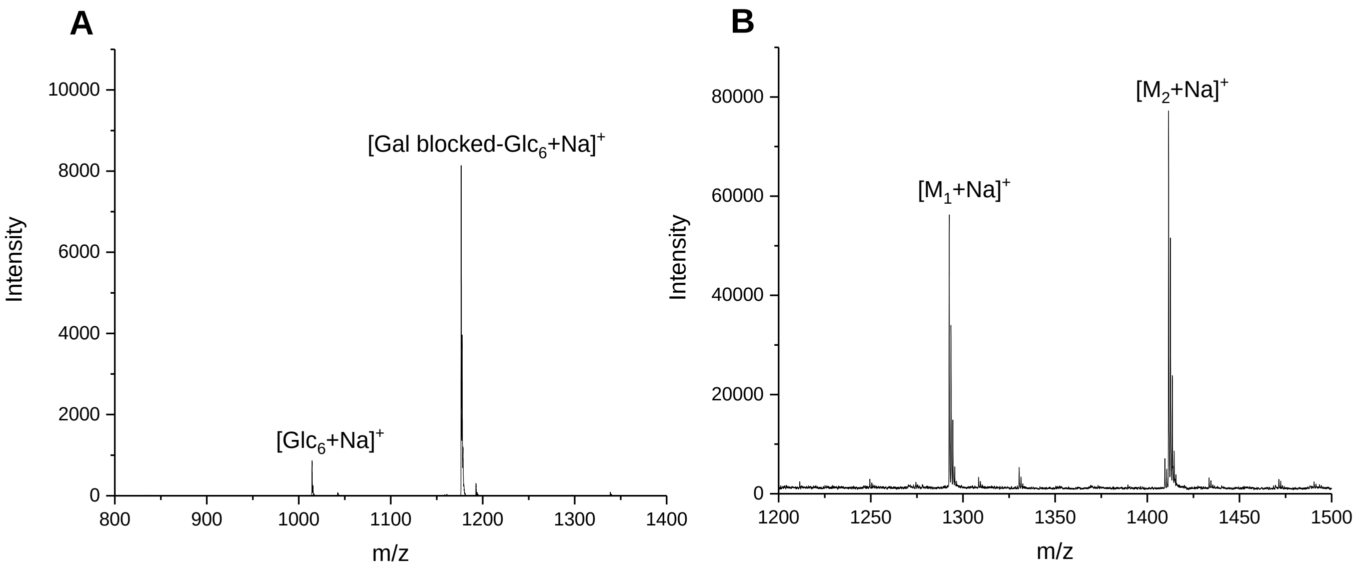

**Supplementary figure 11| Galactosylation and hydrazide ligation of native cellooligosaccharide cellohexaose**

**A** MALDI-TOF MS showing galactosylation of cellohexose by LgtB. **B** MALDI-TOF MS showing sodiated adducts of single ($M_1$ = 1,270) and double ($M_2$ = 1,389) ligation of nicotinic hydrazide to the oxidized, galactose blocked cellohexaose.

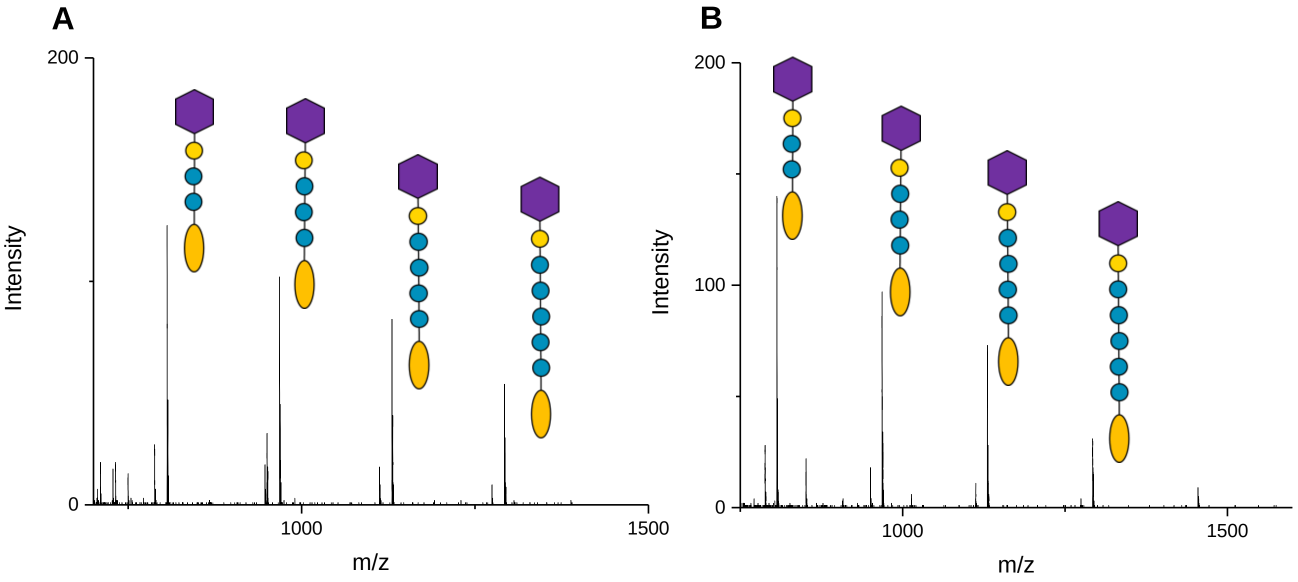

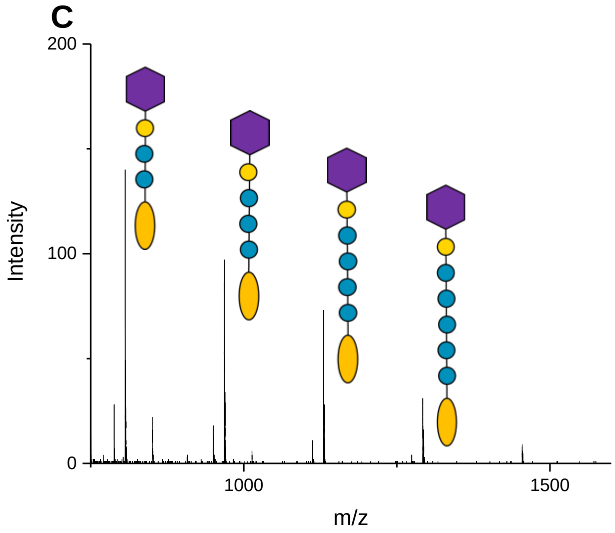

**Supplementary figure 12| Confirmation of β-galactosidase activity blocking**
MALDI-TOF MS showing nicotinic hydrazide ligated ITag-glucose oligosaccharides after incubation with β-galactosidases from glycoside hydrolase families **A** GH1 **B** GH42 and **C** GH50. Nicotinic hydrazide group represented by purple hexagon.

**Supplementary Table 1 | LgtB activity on donor and acceptor substrates as detected by MALDI-TOF MS . (+) observed activity, (-) no activity observed.**

| Acceptors | Donors | | | | |
|---|---|---|---|---|---|
| | UDP-Glc | UDP-Gal | UDP-Xyl | UDP-GalNAc | UDP-GlcNAc |
| Glc-Itag | + | + | + | + | - |
| GlcNac-I-Tag | + | + | + | + | - |
| Cellobiose (Glc-β1,4-Glc) | + | + | | + | - |
| Glc | + | + | | | |
| GlcNAc | + | + | | | |
| GlcN | + | + | | | |
| β-GlcNAc-N3 | + | + | | | |
| β-Glc-N3 | + | + | | | |
| Glc-pNP | + | + | | | |
| Trehalose (Glc-α1,1-Glc) | - | - | | | |
| Man | + | + | | | |
| Sucrose (Glc-α1,2β-Fru) | + | + | | | |
| Cellotetraose-pNP | + | + | | | |
| Lactose (Gal-β1,4-Glc) | - | - | | | |
| Gal | - | - | | | |
| Xyl | - | - | | | |
| Ara | - | - | | | |

### Materials and methods

##### Galactosyltransferase production

Galactosyltransferases LgtB (Uniprot: Q51116), LgtC (Uniprot: A0A3S5C3F9), LgtH (Uniprot: Q2TIJ3), B4GALT4 (Uniprot: B2RAZ5) and an uncharacterized homolog (Uniprot: G0PH97) were provided by Prozomix Lld, Haltwhistle, UK as purified protein suspended in $(NH_4)_2SO_4$. Samples were centrifuged (20000 g, 5 min, 4 °C), the supernatant was discarded and an equal volume of Tris buffer (20 mM pH 8.0) was added to resuspend the protein pellet. The protein sample was centrifuged again to remove precipitated protein and supernatant retained for analysis.

##### NMR of carbohydrates

NMR characterisation of I-Tagged glucose oligosaccharide linkages were performed using a $^{1}H$-$^{13}C$ gradient-selected sensitivity-enhanced multiplicity-edited HSQC on a Bruker AVIII 500 MHz spectrometer equipped with a QCI-F cryoprobe. Assignments were made based on the carbohydrate structure database (http://csdb.glycoscience.ru/database/).

##### Mass spectrometry of carbohydrates

Samples were prepared for analysis by MALDI-TOF via crystallisation with super DHB or THAP matrices. Super DHB (a 9:1 (w/w) mixture of 2,5-Dihydroxybenzoic acid and 2-hydroxy-5-methoxybenzoic acid) was prepared at 15 mg $ml^{-1}$ in a mixture of 50 % (v/v) acetonitrile and 50 % (v/v) water containing 0.1 % trifluoroacetic acid (TFA). 2′,4′,6′-Trihydroxyacetophenone monohydrate (THAP) was prepared at 10 mg $ml^{-1}$ in acetone. MALDI-TOF mass spectrometry was performed using the Bruker Ultraflex 3 in positive mode. MALDI-TOF was calibrated using peptide Calibration Standard II (Bruker) containing a range of peptides 757-3149 Da in size.

##### SuSy system for UDP recycling

*Solanum lycopersicum* Sucrose synthase (SLSUS6) was expressed in *E. coli* BL21(DE3). Cells were cultured at 37 ºC, 250 rpm until reaching an OD600 of 0.6, after which cultures where moved to 18 ºC and protein expression was induced via addition of 1 mM IPTG for 20 h. Cells were harvested and lysed, after which cleared cell lysates were loaded onto an immobilized metal affinity chromatography (IMAC, GE Healthcare) $Ni^{2+}$-charged column. After washing 30 column volumes with 50 mM Tris, 50 mM NaCl, pH 8.0, SLSUS6 was eluted via addition of 500 mM imidazole. SuSy UDP-sugar regeneration reactions were carried out in a total volume of 50 µl containing 50 mM MES buffer (pH 6.0), 0.5 mM I-Tag substrate, 0.5 mM UDP, 35 mM sucrose, 5 mM $MgCl_2$, 2.8 µg SuSy and 20 µg LgtB. Reactions were incubated at 37 $^{o}C$ for 5 days and analysed by MALDI-TOF.

**GalT screening panel**

The panel of galactosyltransferases was screened in reactions as follows: 0.5 mM ITag acceptor, 1.5 mM UDP sugar donor, 10 mM $MgCl_2$, 10 mM $MnCl_2$, 0.1 mg $ml^{-1}$ BSA, 50 mM Tris pH 8.0, and enzyme concentrations, one of the following: LgtH (0.3 mg $ml^{-1}$), G0PH97 (0.4 mg $ml^{-1}$), LgtC (0.1 mg/mL) and LgtB (1.7 mg $ml^{-1}$). Reactions were incubated at 37 ºC for 7 days to allow any potential polymerization to occur. ITag acceptors ITag-Glc, ITag-GlcNAc and ITag-LacNac were utilized in the screen along with UDP donors UDP-Glc and UDP-Gal.

**Donor scope assessment and I-Tagged oligosaccharide production using LgtB**

Final concentrations in I-Tag-glucose oligosaccharides experiments: 15 mM UDP-Glc, 1 mM I-Tag-glucose, 50 mM Tris HCl (pH 8.0), 10 mM $MgCl_2$, 10mM $MnCl_{2,}$ BSA 100 µg $ml^{-1}$, ~1.7 mg $ml^{-1}$ LgtB (resuspended in $dH_2O$). Experiments assessing the activity of UDP-Xyl transfer onto ITag-glucose were as previous but with UDP-xylose 1mM final concentration, replacing UDP-glucose as the donor. I-Tag-glucose oligosaccharides were purified on a C18 columns using a gradient of methanol concentrations.

**Enzymatic digestions of ITagged-oligosaccharides**

β-glucosidase from almond (49290, Sigma-Aldrich) was dissolved in water at 1 mg $ml^{-1}$. Cellulase from *Aspergillus niger* (C1184**,** Sigma-Aldrich) was dissolved in pH 5.0, sodium acetate buffer at 10 mg $ml^{-1}$. Reactions were performed at pH 5.0, 37 °C for 2 h at concentrations of 0.5 mg $ml^{-1}$ and 5 mg $ml^{-1}$ β-glucosidase and cellulose respectively with 10 % (v/v) C18 column purified I-Tagged glucose oligosaccharides. LPMO reactions were set up as follows: 100 µM $H_2O_2$, 200 µM ascorbate and 5 µM LPMO in 25 mM Tris-HCl buffer pH 7.5 and performed at room temperature for 14 h.

**LPMO production**

The genes coding for *Aspergillus oryzae* AoLPMO11 (Uniprot: Q2UA85), *Neurospora crassa* NcLPMO9C (Uniprot: Q7SHI8) and *Thermobifida fusca Tf(*AA10)B (Uniprot: Q47PB9) cloned into pET22b and expressed using *E. coli* C43(DE3), via periplasmic secretion to obtain an N-terminal His. After growth at 37°C until OD600 of 0.6, protein expression was induced with 0.1mM IPTG at 25°C for 16 h, 200 rpm. *Lentinus similis* LsLPMO9C (Uniprot: A0A0S2GKZ1) was expressed as an E8K-vector construct in *E. coli DH5a* at 25 °C for 24 h, 200 rpm. Protein production was induced using 10mM arabinose.

Cells were lysed in equilibration buffer (50 mM sodium phosphate (NaPi) buffer pH 8.0, containing 300 mM NaCl, 1 mg $ml^{-1}$ lysozyme and 10 µg $ml^{-1}$ DNase,) via sonication, after which cleared supernatant was applied to a pre-equilibrated 5 ml Strep-tactin superflow cartridge (Qiagen). The column was washed with equilibration buffer and the protein eluted using 5mM desthiobiotin.

Fractions containing pure target protein, identified via SDS-PAGE, were pooled, concentrated and re-buffered into 25 mM Tris-HCl pH 7.5 using PD10 desalting columns. The protein was loaded with 5x molar excess $CuCl_2$ for 1 h at 4 °C and the excess Copper was then removed using PD10 desalting columns. The protein was stored at -80 °C, 25 mM Tris-HCl buffer pH 7.5.

**Endo-substrate development**

$M_1$ Galactose oxidase (GOase) was expressed and purified as previously described[1]. Oxidized UDP-Gal (UDP-$Gal_{Ox}$) was produced as follows: 0.1 mg $ml^{-1}$ horseradish peroxidase (HRP), 1 mg $ml^{-1}$ GOase $M_1$, 0.1 mg $ml^{-1}$ catalase, 10 mM UDP-galactose, NaPi pH 7.4 reaction was incubated at 25 $^{o}$C, 250 rpm for 4 h. Final concentrations in oxidized UDP-Gal blocking of cellohexaose were as follows: 10 mM UDP-$Gal_{Ox}$, 5 mM cellohexaose, 10 mM $MnCl_2$, 50 mM NaPi (pH 8), LgtB (1.7 mg $ml^{-1}$). Final concentrations in $Gal_{Ox}$ blocking of I-tagged glucose oligosaccharides were as follows: 10 mM UDP-$Gal_{Ox}$, 20 % (v/v) C18 column purified I-tagged glucose oligosaccharides, 10 mM $MnCl_2$, 50 mM sodium phosphate (pH 8), LgtB (1.7 mg $ml^{-1}$). After incubation at 250 rpm, 37 $^{\circ}$C for 16 h, reaction products were purified on a viva spin column (10000 MWCO). Subsequent hydrazide ligation required 20 equivalents of nicotinic hydrazide addition to the previous reaction mixes, which were then incubated at 30 $^{\circ}$C, 250 rpm for 2 h.

Enzymatic galactosylation of native cello-oligosaccharides (Fig. S11a) followed by oxidation of the terminal Gal then nicotinic hydrazide ligation (Fig. S11b) was also demonstrated. Final concentrations in galactose capping of cellohexaose were as follows: 10 mM UDP-Gal, 5 mM cellohexaose, 10 mM $MnCl_2$, 50 mM NaPi (pH8), LgtB (1.7 mg $ml^{-1}$). Reaction was incubated at 250 rpm, 37 $^{\circ}$C for 16 h. Reaction was then purified on a viva spin column. Product was then oxidized, final concentration during the oxidation of the galactose capped cellohexaose were as follows: 0.1 mg $ml^{-1}$ HRP, 1 mg $ml^{-1}$ GOase $M_1$, 0.1 mg $ml^{-1}$ catalase, 0.5 mM galactosylated cellohexaose, NaPi pH 7.4. Reaction was incubated at 25 $^{\circ}$C, 250 rpm for 4 h then treated with nicotinic hydrazide as in previous method.

Nicotinic hydrazide ligated I-Tagged cello-oligosaccharides were incubated with β-galactosidases to ensure inaccessibility of exo-active cellulolytic enzymes. Reactions with GH1 were incubated at 90 $^{\circ}$C for 2 h in reactions containing: nicotinic hydrazide ligated I-Tagged cello-oligosaccharides (25 % v/v), 50 mM pH 5 sodium acetate buffer, 0.5 mg $ml^{-1}$ GH1. Reactions with GH42 were incubated at 65 $^{\circ}$C for 2 h, in reactions containing: nicotinic hydrazide ligated I-Tagged cello-oligosaccharides (25 % v/v), 50 mM pH 4 sodium acetate buffer, 0.5 mg $ml^{-1}$ GH42. Reactions with GH50 were incubated at 40 $^{\circ}$C for 2 h, in reactions containing nicotinic hydrazide ligated I-Tagged cello-oligosaccharides (25 % v/v), 50 mM pH 7 Tris buffer, 0.25 mg $ml^{-1}$ GH50.

**Synthesis of 4-(1-Methyl-3-methyleneimidazolium)benzyl β-D-glucopyranoside trifluoromethanesulfonate (2)**

Glycosyl acceptor 4-(chloromethyl)benzyl alcohol (0.0689 g, 0.440 mmol, 1 eq) and glycosyl donor 2,3,4-tri-O-benzoyl-6-O-chloroacetyl-α-D-glucopyranosyl trichloroacetimidate (0.6277 g, 0.880 mmol, 2.0 eq) were placed in a dry vial and dried under vacuum for 30 min. 2.20 ml of anhydrous DCM was added to the donor/acceptor vial under nitrogen, resulting in a solution of volume 2.75 ml and therefore approximately 0.160 M in acceptor and 0.320 M in donor. A stock solution of TMSOTf (0.06 M in DCM) was made by dissolving TMSOTf (0.217 ml, 1.200 mmol) in 20 ml anhydrous DCM. 3.0 ml of this stock solution was used for this reaction. The flow microreactor and attached tubing was flushed with nitrogen. The donor/acceptor solution and TMSOTf solution were each taken up in a syringe and installed onto a syringe pump. The solutions were then injected into the microreactor (total internal volume of reactor chip and outlet tubing = 32.8 μL) at the desired flow rate corresponding to the residence time (15 seconds, 65.60 μl $min^{-1}$ in each syringe for a combined flow rate of 131.20 μl $min^{-1}$ in reactor zone) via the inlet tubing. The flow reaction was performed at RT. The mixture that flowed from the microreactor was dropped in a flask containing reagent grade DCM in air to quench the reaction. Reaction solution was collected for 41 min 30 sec, after which time the reaction mixture solvent was removed under reduced pressure. The crude product was dissolved in DCM (20 ml) and washed with water (8 ml), then the water was extracted with a further portion of DCM (20 ml). The DCM fractions were collected, dried with magnesium sulfate, filtered and the solvent was removed under reduced pressure. The dried residue was dissolved in a minimal volume of DCM, then HCl, 1.25 M in MeOH (7.04 ml, 8.800 mmol) was added. The resulting solution was stirred for 16 h at RT in air, then diluted with DCM (15 ml) and water (10 ml) and product was extracted into the DCM phase. The aqueous phase was washed with a further DCM portion (15 ml) then DCM washings were combined, dried using magnesium sulfate, filtered and the solvent was removed under reduced pressure. The dried residue was washed with hexane (3 x 5 ml), then dried under reduced pressure for 1 h, before being dissolved in anhydrous MeCN (5 ml) under a nitrogen atmosphere. 1-Methyl imidazole (0.14 ml, 1.76 mmol) and potassium trifluoromethanesulfonate (0.3312 g, 1.76 mmol) were added and the resulting mixture was heated under reflux at 90 ˚C and stirred for 18 h, after which time TLC (DCM:MeOH 94:6) showed the reaction to be complete by MS. Solvent was removed under reduced pressure, then DCM (5 ml) and 1 M HCl $_{aq.}$ (5 ml) were added to the residue and product was extracted into the DCM phase. The aqueous phase was washed with DCM (2 x 5 ml), then the DCM portions were combined, dried using magnesium sulfate, filtered and the solvent was removed under reduced pressure. The crude residue was washed with neat $Et_2O$ (5 ml) and DCM:$Et_2O$ 5:95 (2 x 5 ml). The I-Tagged products were dissolved in methanol (2.2 ml) and sodium methoxide (50.0 μl, 0.22 mmol, 25 % wt in MeOH) was added. The solution was stirred at RT for 3 h after which time TLC-MS showed the reaction to be complete. The solution was then brought to pH 7 using 1 M HCl $_{aq.}$. Solvent was removed, then the residue was diluted with DCM and water and product was extracted into the aqueous phase. Water was removed under reduced pressure and the dried mixture was purified by reverse-phase HPLC (Water:MeCN) to yield the **I-Tag-glucose 2 (2)** (0.1110 g, 49 % over 4 steps) as a solid.

**$^1$H NMR** $\delta_H$ (500 MHz, Methanol-$d_4$) for ITag-glucose **2**

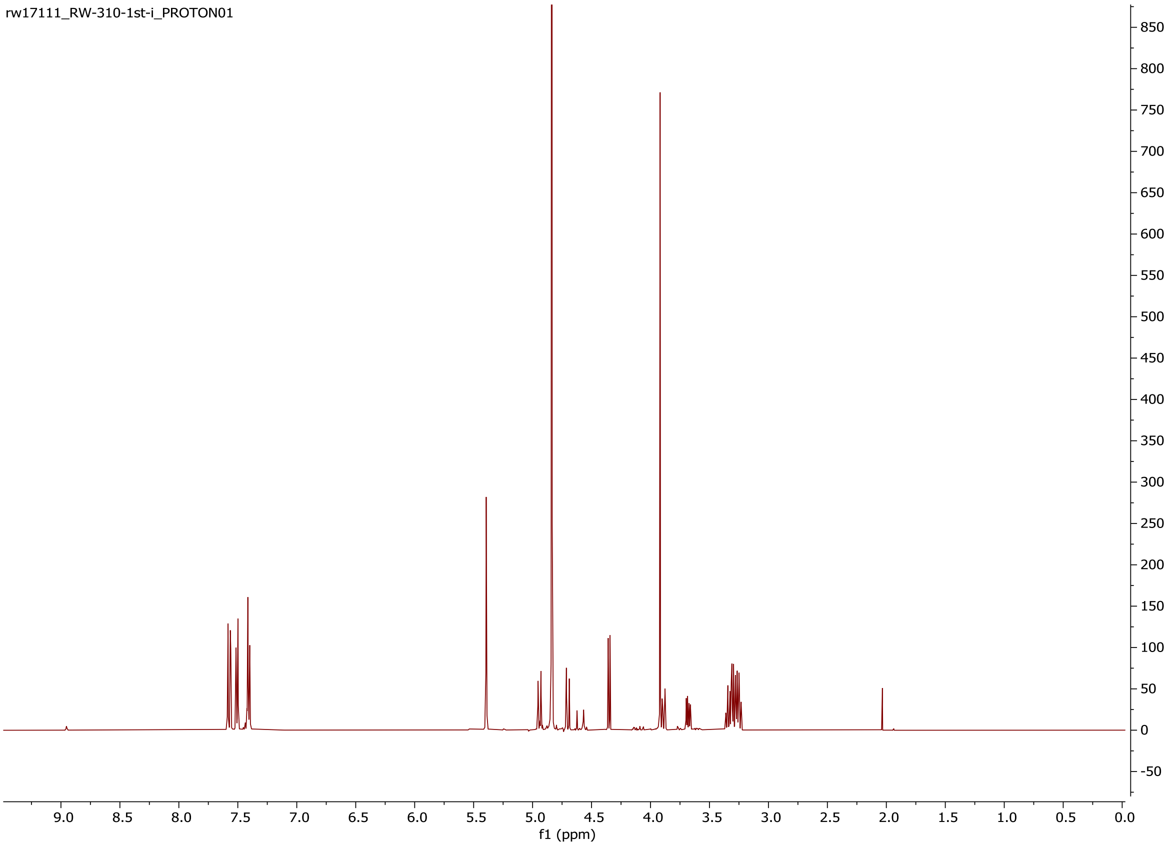

**$^{13}$C NMR** δ $_C$ (126 MHz, Methanol-$d_4$) for ITag-glucose **2**

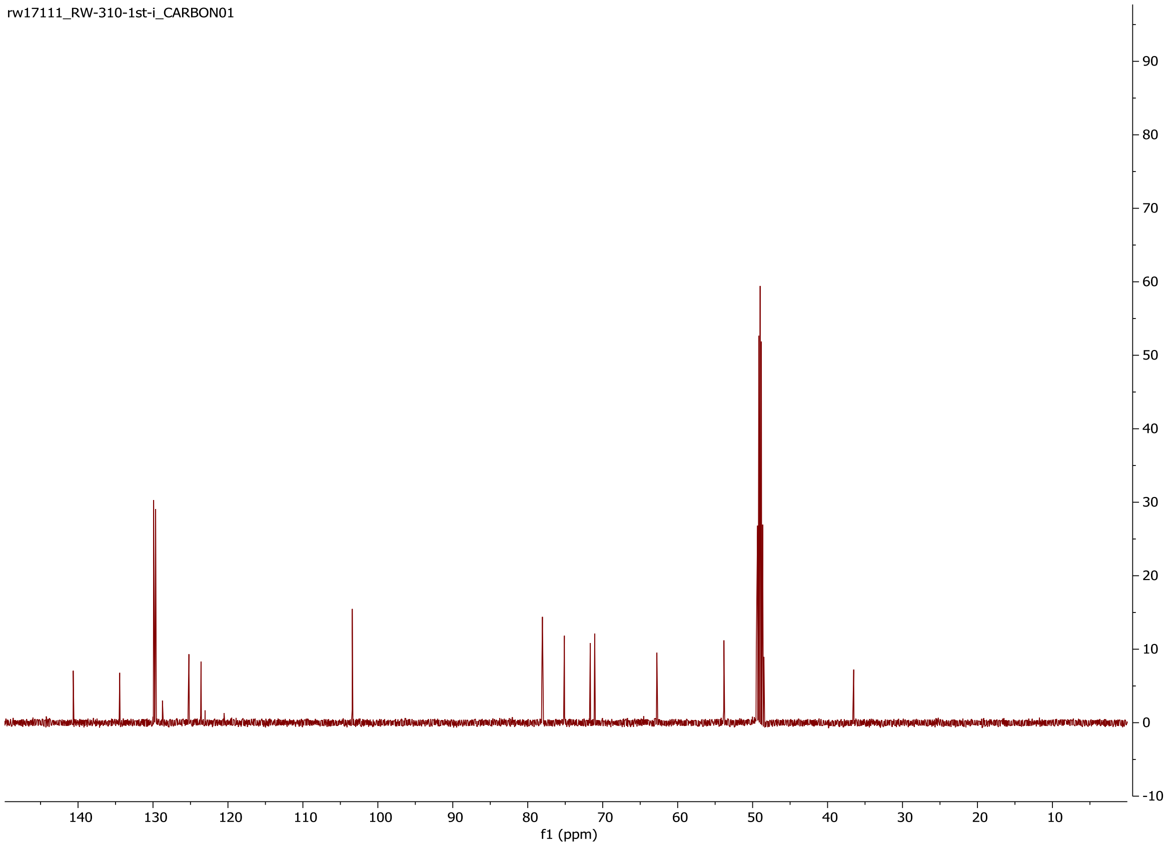

(1) Herter, S.; McKenna, S. M.; Frazer, A. R.; Leimkühler, S.; Carnell, A. J.; Turner, N. J. Galactose Oxidase Variants for the Oxidation of Amino Alcohols in Enzyme Cascade Synthesis. *ChemCatChem* **2015**, *7* (15), 2313–2317.
